# Decision by sampling implements efficient coding of psychoeconomic functions

**DOI:** 10.1101/220277

**Authors:** Rahul Bhui, Samuel J. Gershman

## Abstract

The theory of decision by sampling (DbS) proposes that an attribute’s subjective value is its rank within a sample of attribute values retrieved from memory. This can account for instances of context dependence beyond the reach of classic theories which assume stable preferences. In this paper, we provide a normative justification for DbS that is based on the principle of efficient coding. The efficient representation of information in a noiseless communication channel is characterized by a uniform response distribution, which the rank transformation implements. However, cognitive limitations imply that decision samples are finite, introducing noise. Efficient coding in a noisy channel requires smoothing of the signal, a principle that leads to a new generalization of DbS. This generalization is closely connected to range-frequency theory, and helps descriptively account for a wider set of behavioral observations, such as how context sensitivity varies with the number of available response categories.

---

Descriptive accounts of decision making, such as expected utility theory and prospect theory (Kahneman & Tversky, 1979), are typically based on a stable set of “psychoeconomic” functions specifying the mental representations of gains, losses, probabilities, and delays. However, the psychological reality of such functions has been challenged by evidence that decisions are highly context-sensitive: the mental representation of an attribute changes depending on the choice set and other attribute values retrieved from memory (Vlaev, Chater, Stewart, & Brown, 2011). As an alternative, some accounts have proposed that decisions are based on more elementary cognitive operations, namely memory retrieval and comparison (Johnson, Haubl, & Keinan, 2007; Marchiori, Di Guida, & Erev, 2015; Stewart, 2009; Stewart, Chater, & Brown, 2006). One particularly influential account*—decision by sampling* (DbS; Stewart, 2009; Stewart et al., 2006)—attempts to reconcile these viewpoints by proposing that psychoeconomic functions can be derived from principles of memory retrieval and comparison. According to DbS, the shapes of these functions are malleable, reflecting both local context and long-term statistical regularities that constitute the database from which memories are sampled. Despite its simplicity, DbS has accounted for a wide range of empirical phenomena (Stewart, Chater, Stott, & Reimers, 2003; Stewart, Reimers, & Harris, 2014; Ungemach, Stewart, & Reimers, 2011; Walasek & Stewart, 2015).

The basic idea of DbS is that attributes are sampled from memory and ordinally compared to the attributes of the current prospect. By tallying these ordinal comparisons, a decision maker computes the rank of the prospect’s attribute relative to the distribution of attribute magnitudes in memory. Preference is then determined by comparing the ranks across prospects. While DbS is a psychological process model, this paper shows that the same set of ideas can be arrived at through a normative analysis. In particular, we derive DbS from the principle of *efficient coding,* which has a long history in the study of perceptual systems (Atick & Redlich, 1992; Attneave, 1954; Barlow, 1961; Laughlin, 1981), and has more recently been applied to value representation in the brain (Louie, Grattan, & Glimcher, 2011; Louie, Khaw, & Glimcher, 2013; Louie, LoFaro, Webb, & Glimcher, 2014; Rangel & Clithero, 2012).

According to the efficient coding principle, the brain is designed to communicate information with as few spikes as possible, since spikes are metabolically expensive. As will be described in more detail below, this is accomplished by choosing a neural code that maximizes the mutual information between a neuron’s inputs and outputs. When neurons are conceived as noiseless communication channels, maximizing mutual information is equivalent to minimizing redundancy, which can be achieved by recoding inputs according to their rank—precisely the operation implemented by DbS in the limit of an infinite number of samples. However, the channel becomes noisy when only a finite number of samples are drawn from memory, in which case some redundancy in the code is required to suppress noise. An approximation of the information-maximizing strategy is to smooth the samples prior to the rank transformation. This leads to modifications of DbS that were previously proposed to account for range effects on choice (Brown & Matthews, 2011; Parducci, 1995; Ronayne & Brown, 2017; Stewart et al., 2006).

The central contribution of our work is to clarify the computational design principles of DbS and related models, uniting them with an important strand of theoretical neuroscience. This paves the way for new behavioral predictions, insights into how DbS might be implemented in the brain, and a deeper understanding of the connections between information theory and decision making.

## Decision by sampling

In this section, we present DbS formally and then review its applications to empirical phenomena.

Let *x* ∈ ℝ denote an attribute value in a psychoeconomic space (e.g., gains, losses, probabilities, delays). This attribute occurs in the environment with probability distribution *f*(*x*).

DbS samples a set of comparison values x_1:*N*_ = {*x_1_,…, x_N_*} from *f*(*x*) and then computes the rank of *x* relative to the sample:

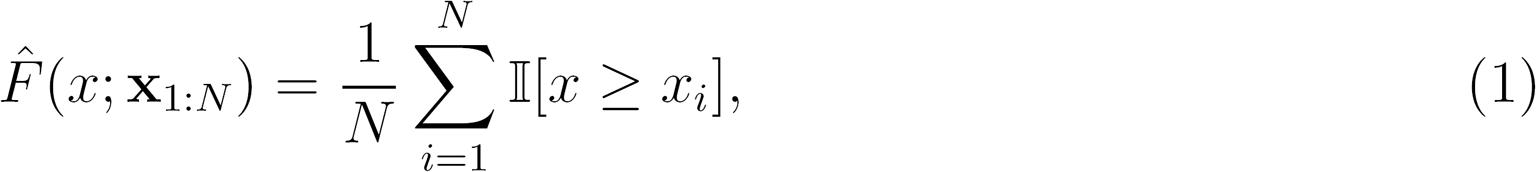

where Ⅱ[‧] = 1 when its argument is true, 0 otherwise.^1^ In the infinite sample limit, the rank function converges with probability 1 to the cumulative distribution function (CDF), F(*x*):

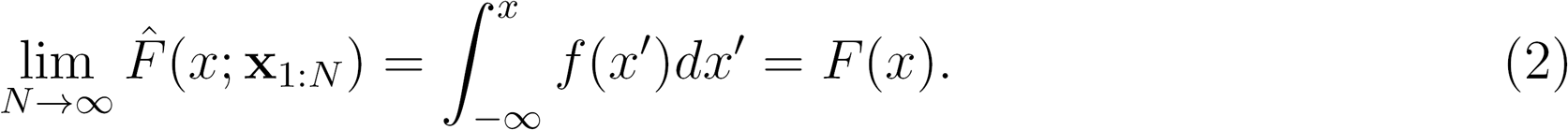

The rank function (and the CDF) is monotonic in *x*, but importantly it exhibits steeper changes in high probability regions of the attribute space. Stewart et al. (2006) used this property to explain several well-known properties of psychoeconomic functions, using various proxy estimates of *f*(*x*) for different attributes:

- Small gains and losses are more probable than large gains and losses, thus the rank functions for gains and losses are concave (diminishing marginal utility). Importantly, small losses are relatively more likely than small gains, implying that the rank function for losses is steeper, as proposed in prospect theory (Kahneman & Tversky, 1979).
- The distribution of temporal delays approximately follows a power law, giving rise to a power-law rank function. This subsumes hyperbolic discounting as a special case, but in fact the best-fitting rank function is sub-hyperbolic, consistent with several experimental studies (Myerson & Green, 1995; Simpson & Vuchinich, 2000).
- Very small and very large probabilities are more commonly encountered than midrange probabilities, giving rise to an inverse S-shaped rank function (i.e., overweighting of low probabilities and underweighting of high probabilities), in agreement with the probability weighting function derived from choice experiments (Gonzalez & Wu, 1999) and postulated by prospect theory.

One of the principal advantages of DbS over traditional approaches is that it can explain departures from these properties as the result of transient contextual information that distorts the long-term statistical regularities. Stewart et al. (2014) showed that the shapes of utility, discount, and probability weighting functions could be altered by exposing subjects to distributions of attribute values that varied in their skew. For example, a concave utility function could be converted to a convex function simply by populating the set of large gains more densely than the set of small gains, and the rate of discounting could be slowed by sampling delays from a uniform distribution (rather than an ecologically valid distribution with positive skew). Field studies by Ungemach et al. (2011) have recapitulated these observations, finding that choices between two lotteries were affected by incidental exposure to intermediate attribute values (supermarket prices), and choices between two delayed outcomes were affected by exposure to events occurring at intermediate delays. Large-scale studies of satisfaction as a function of income make the same point: relative income rank strongly determines satisfaction (Boyce, Brown, & Moore, 2010; Brown, Gardner, Oswald, & Qian, 2008).

## An efficient coding perspective

What is the computational logic of the rank transformation? To shed some light on this question, let us view psychoeconomic functions as *communication channels,* taking as input an attribute value *x* and emitting as output a signal *y* drawn from the probability distribution *f*(*y*|*x*). In designing such a channel, a basic problem is that the amount of information that can be reliably transmitted over a channel with fixed transmission rate (the channel capacity) is finite (Shannon & Weaver, 1949). A neuron consumes several orders of magnitude more energy during spiking compared to rest, such that the brain’s energy budget can only afford to have around 1% of neurons active at any time (Lennie, 2003). Thus, the energy budget imposes a stringent constraint on the channel capacity of neurons, placing demands on neural codes to communicate information with as few spikes as possible. Consistent with this proposition, studies of many different neural systems suggest that economizing on spikes is a fundamental design principle (Laughlin, 2001).

There are two strategies to reduce the cost of information transmission. One is to reduce the signal-to-noise ratio (i.e., transmit lower precision messages); we will not consider this strategy further here, under the assumption that organisms need to maintain a certain level of precision for survival. The second strategy is to eliminate redundancy by recoding inputs. Intuitively, if an input can be predicted before the output has been observed, then the output is not conveying any information about the input—it is redundant with the receiver’s prior knowledge. In other words, unpredictable outputs are more informative than predictable outputs. The most unpredictable output distribution is uniform; hence, the goal of redundancy reduction is to find a code that gets the output distribution close to uniform.

As a rough illustration, suppose one wants to identify an item value which can take on four possible levels with equal probability: very low, slightly low, slightly high, or very high. Consider the binary code shown in Figure 1a which represents “very low” value as the *codeword* 00, “slightly low” as 01, “slightly high” as 10, and “very high” as 11 (assume no mistakes occur in this encoding process). All codewords are of length 2 which means that each value can be identified by 2 binary digits (aka *bits*). Each bit can intuitively be thought to capture a yes-or-no question that helps to distinguish some values from others. For example, in this code, one bit reflects the question “Is the value high or low?” while the other reflects the question “Is the value slightly or very high/low?”

In biological terms, ones and zeros may be expressed as different configurations of neuronal activity. However, organisms have a finite number of neurons, each with bounded precision and response range, and limited metabolic energy to spend on signaling. An accurate representation of the world demands efficient use of these constrained resources. Roughly speaking, a neural code is considered to be more efficient if it requires fewer bits on average to identify an input. If the input distribution is not uniform, efficient codes prioritize values that occur more frequently by assigning them shorter codewords. Thus the more efficient code in Figure 1b expends one full bit to pinpoint the “very low” value level, which is the most common. Its expected codeword length is 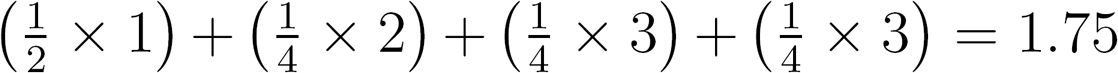 bits, making it superior to the fixed-length 2-bit code (and in fact, it reaches peak efficiency in this situation).

This efficiency obtains because, for the non-uniform attribute distribution, asking whether the value is low (versus high) overlaps substantially with asking whether the value is very low (versus slightly low). Most of the time, an affirmative answer to the first query is accompanied by an affirmative answer to the second. Hence the less efficient code, which is based on these queries, exhibits redundancy.^2^ A better approach is to consolidate these questions and ask straight away whether the value is very low (versus any other value). Analogous to the game of 20 questions, good codes (or questions) are those that partition the space of possibilities equally. A direct consequence of this strategy is that the output states become unpredictable, reflecting high information gain from each question. Accordingly, notice that the overall distribution of ones and zeros generated by the (more) efficient code is uniform, compared to the less efficient code which outputs more than twice as many zeros as ones. This unpredictability is the hallmark of redundancy minimization in the information-theoretic framework.

To make these ideas more formal, we consider the problem in which an attribute magnitude *x* is drawn from a continuous distribution with CDF *F*(*x*), and must be encoded by an internal representation *y* that takes on one of *M* + 1 possible integral values rescaled to the unit interval, 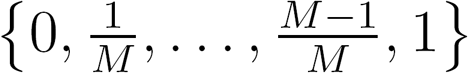. The mutual information between stimulus *x* and response *y* describes the information that *x* and *y* carry about each other, and is defined as

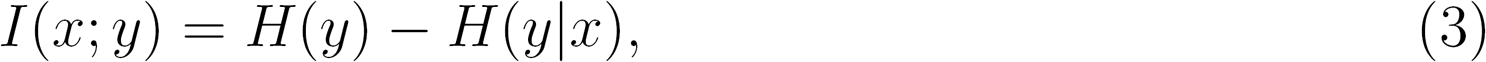

where *H*(*y*) is the entropy of the response and *H*(*y*|*x*) is the conditional entropy of the response given the stimulus (Cover & Thomas, 2006). Entropy in bits is approximately the number of yes-or-no questions required on average to identify stimuli. Noise in the channel is captured by *H*(*y*|*x*) which reflects the residual uncertainty in the response, knowing the stimulus; it expresses how many bits are lost on average when transmitting *x* over the channel. Mutual information can equivalently be written as *H*(*x*) — *H*(*x*|*y*), which clarifies its quantitative interpretation: it is the expected reduction in codeword length (i.e., the number of bits) needed to identify *x* after observing its representation y. As typically applied, the principle of efficient coding entails that stimuli should be encoded to maximize mutual information—that is, the mapping from *x* to *y* should maximize *I*(*x*; *y*).

**Figure 1:**
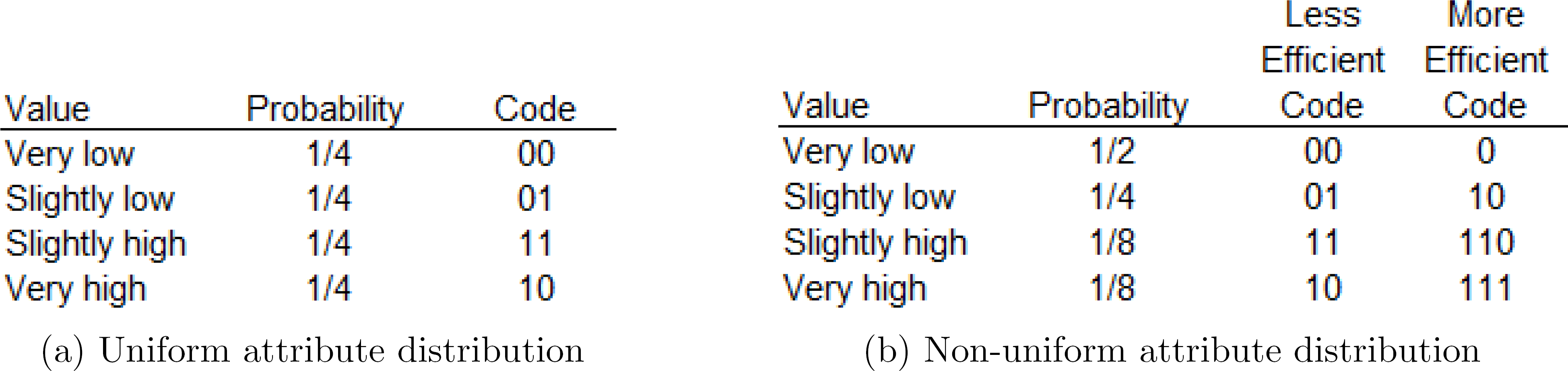
Efficient codes prioritize highly probable regions of the stimulus distribution.

In the noiseless regime, *H*(*y*|*x*) is 0, so maximizing mutual information is equivalent to maximizing output entropy (i.e., unpredictability). Because the maximum entropy distribution for a discrete variable is the uniform distribution, this is achieved by encoding *x* as a discretized version of its CDF, such that the boundaries between representational states arise from the distribution’s (*M* + 1)-quantiles. When the representational states are vanishingly small (i.e., *M* → ∞), this strategy corresponds to the code *y* = *F*(*x*), also known as the *probability integral transform* (Laughlin, 1981). This solution is unique among monotonically increasing response functions. Intuitively, the efficient code prioritizes distinctions between frequently-occurring stimulus values, which are located in regions where the PDF is high and hence the CDF (the integral of the PDF) is steep.

In practice, the brain may not have direct access to the exact distribution function needed to implement this coding scheme. However, samples from the distribution may be available, such as items retrieved from memory. Potential codes then involve (possibly stochastic) mappings from the representations distinguishable by the sample, 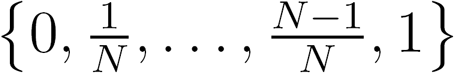, to the available response states, 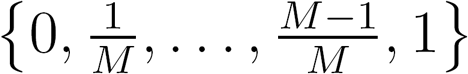. When the sample size is large, the (discretized) empirical rank 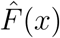 serves as a good approximation of the (discretized) true rank 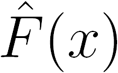. Since DbS approximates the probability integral transform in this way, it can be understood as implementing efficient coding of psychoeconomic functions. In other words, DbS removes redundancies from the representations of gains, losses, probabilities, and delays, so that they can be represented with fewer bits (and thus presumably a lower metabolic cost).^3^

## Coding with finite samples

We can extend this perspective by considering the noisy regime, in which case the resulting code will be corrupted and the probability integral transform is no longer optimal. In particular, when the sample is finite—which is necessarily the case under the inherent computational constraints that organisms face—the empirical rank estimate suffers. Then the optimal code will have some redundancy in order to prevent information loss.

From an information-theoretic perspective, finite samples cause two issues, which we call the *restricted resolution problem* and the *sampling variability problem.* To illustrate, suppose the representational state space is rich (i.e., *M* is large) but only a single sample is drawn (i.e., *N* =1). The quantile-based representation can only take on two possible values in this situation: rank 0 if the sample is higher than the target or rank 1 if the sample is lower. Thus, even though *M* + 1 states are available, the representation is effectively restricted to just two of them (or more generally, *N* +1). As a result, *y* cannot be uniformly distributed across all possible representations, so *H*(*y*) declines; this is the restricted resolution problem. Furthermore, the (*M* + 1)-quantiles that define the code are determined from the sample, and since the sample is stochastic, the representational mapping is noisy. The conditional distribution of *y* is no longer close to a deterministic function of *x*, so *H*(*y*|*x*) rises; this is the sampling variability problem. These issues prevent mutual information from reaching its theoretical maximum of *I*(*x*; *y*) = log(*M* + 1) attained by a perfectly uniform output distribution and a noiseless channel.

The impact of these problems can be quantified more precisely. Continuing to suppose that *M* > *N*, observe that the conditional distribution of *y* given *x* is a rescaled binomial distribution. The number of trials is equal to the sample size *N*, and the probability of success is equal to the probability that a draw is less than *x*—that is, *F*(*x*). Then the marginal distribution of *y* can be derived by integrating its conditional distribution over *f*(*x*):

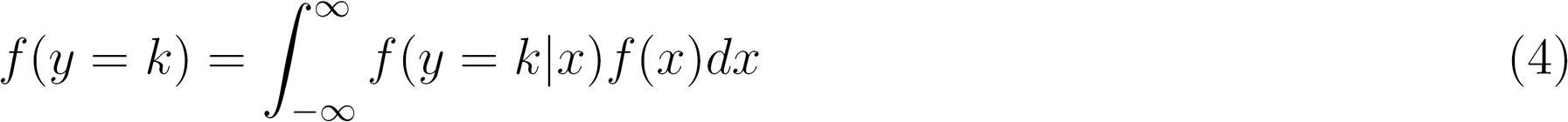

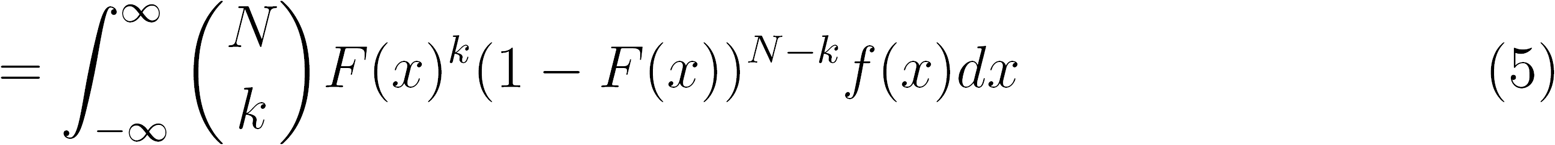

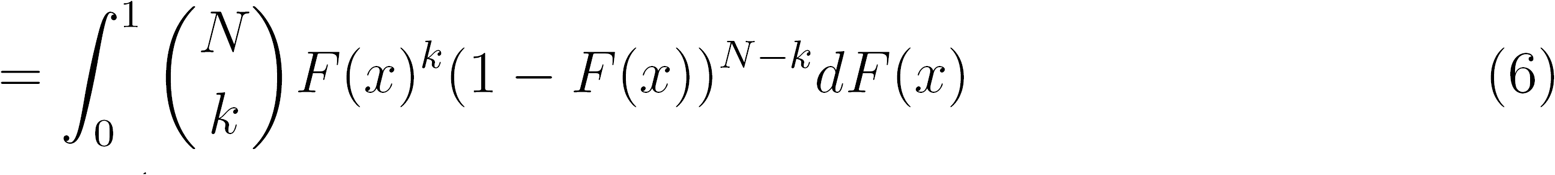

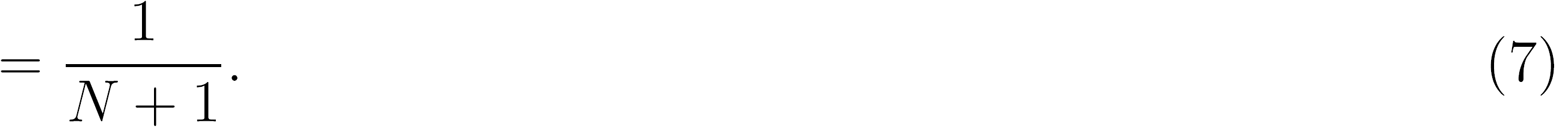

Thus, the output distribution is uniform across the *N* +1 states distinguishable by the sample, and its entropy is

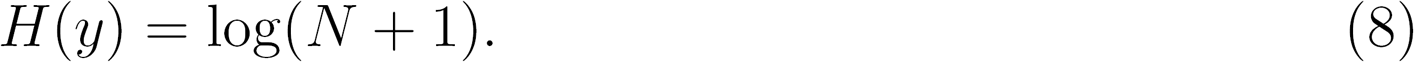

The conditional entropy *H*(*y*|*x*) is more challenging to calculate. A clean exact solution is feasible when *N* = 1, in which case the conditional distribution reduces to a Bernoulli distribution, which has the binary entropy function *H_b_(p)* = —*plogp* — (1 — *p*) *log*(1 — *p*). Substituting *p* = *F*(*x*) and integrating this function with respect to the distribution of *x* yields the conditional entropy:

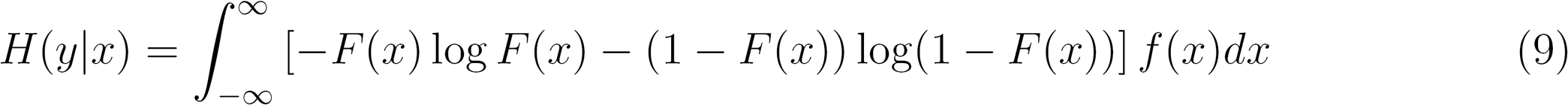

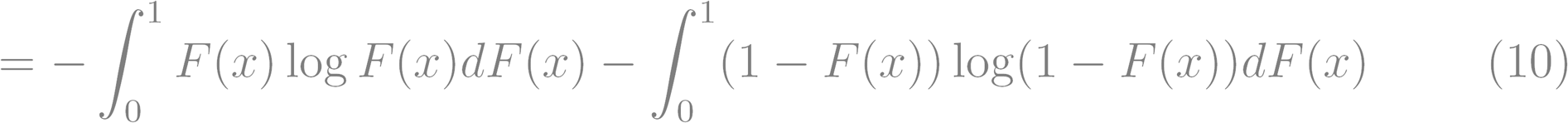

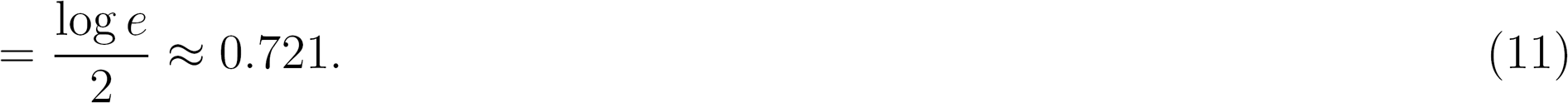

Therefore the mutual information is 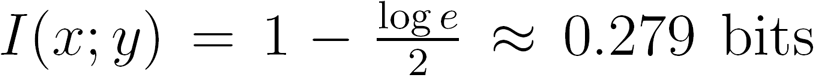; sampling variability costs most of the capacity remaining in the restricted representation. More generally, the entropy of the binomial distribution is known to be approximately 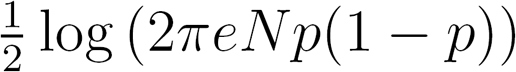. The conditional entropy can thus be approximated by integrating this function over the distribution of *x* after substituting *p* = *F*(*x*):

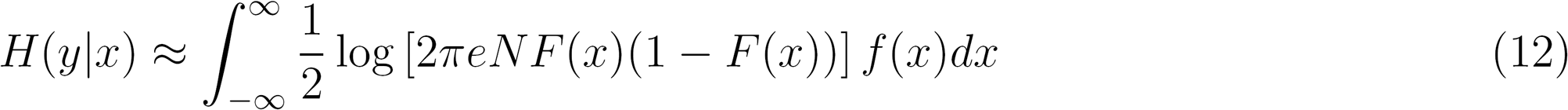

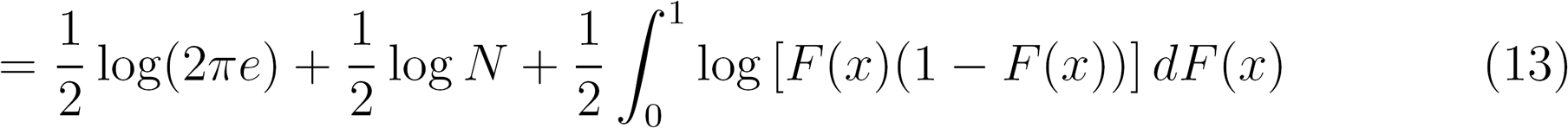

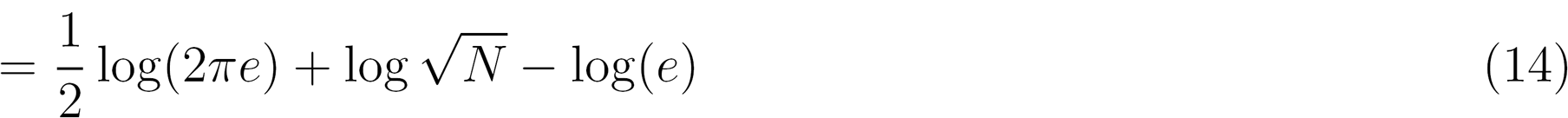

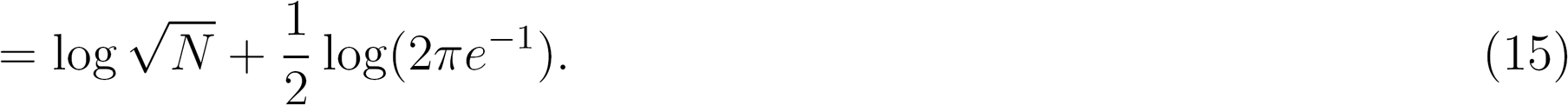

**Figure 2:**
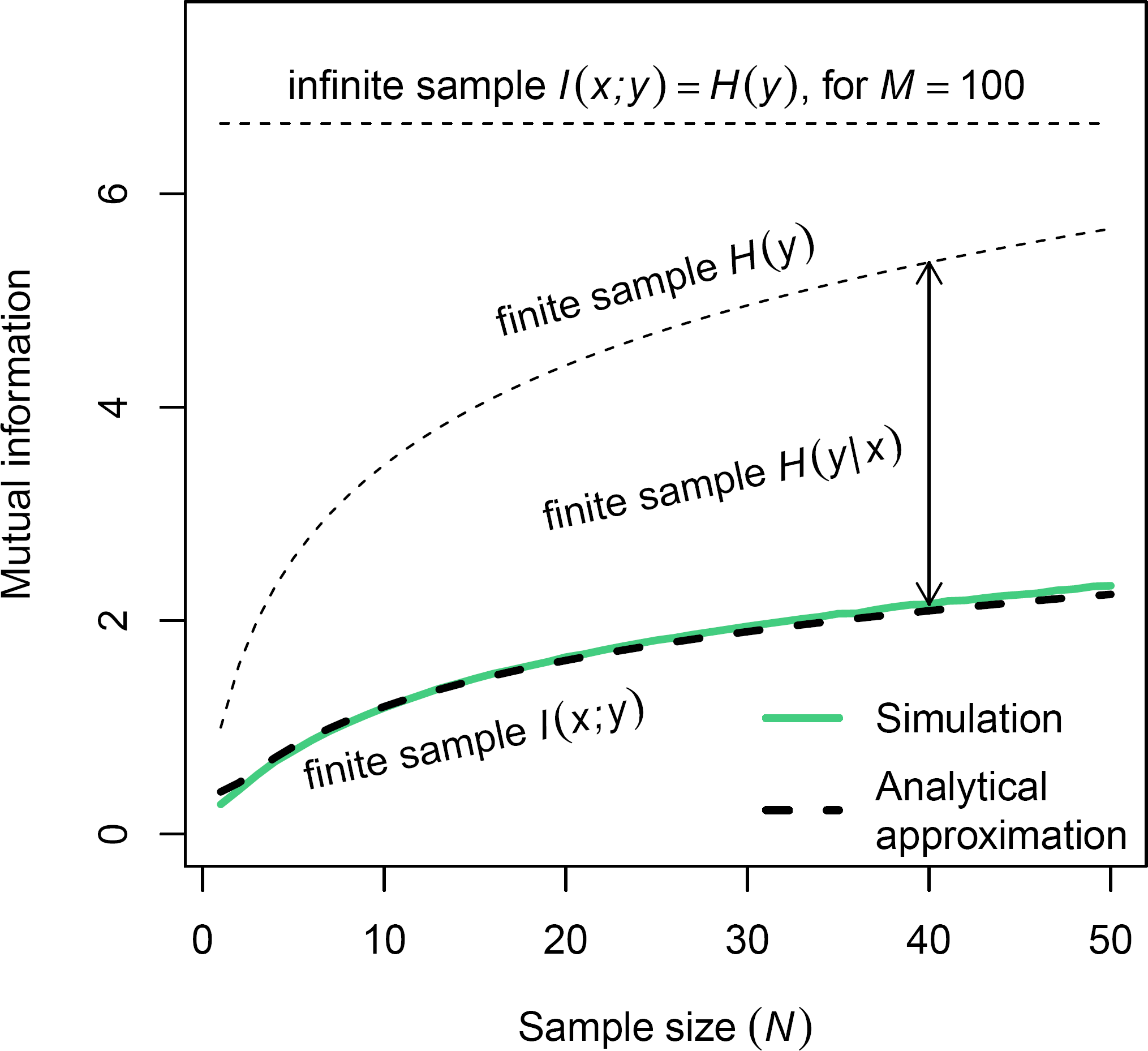
Finite samples reduce output entropy *H*(*y*) and increase conditional entropy *H*(*y*|*x*), causing a decline in mutual information I(x; y). Thick lines represent mutual information in bits between *x* and *y* (both discretized with bin width Yiooo in the simulation). Attribute values *x* are drawn from a *Beta*(1, 2) distribution.

Hence the mutual information is approximately

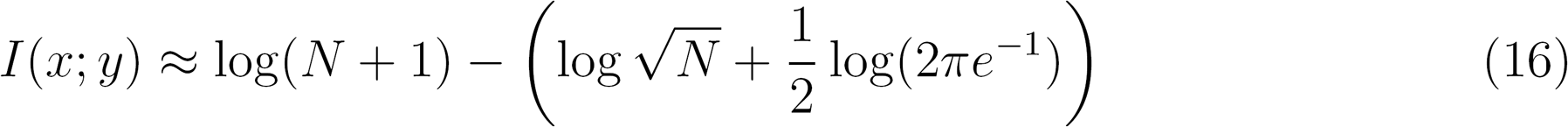

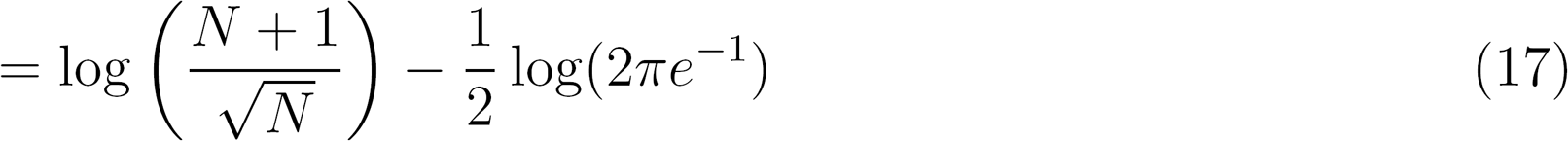

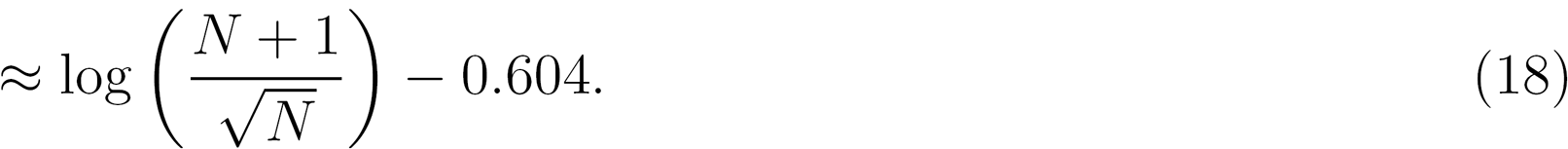

When *N* is large, 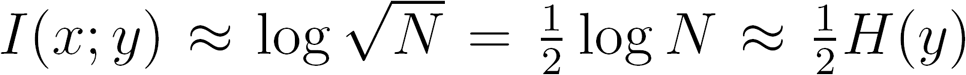. Thus, as a rough rule of thumb, sampling variability chops the restricted capacity in half. Note that all of the above results obtain without specific assumptions about the distribution of *x*. Figure 2 displays the costs imposed by finite samples on mutual information. The values from a simulation closely match the analytical approximation in Equation (18).^4^

One can heuristically satisfy the conflicting demands of redundancy reduction and information transmission by first smoothing the inputs prior to computing the probability integral transform (Atick & Redlich, 1992), such that the response is a discrete analogue of

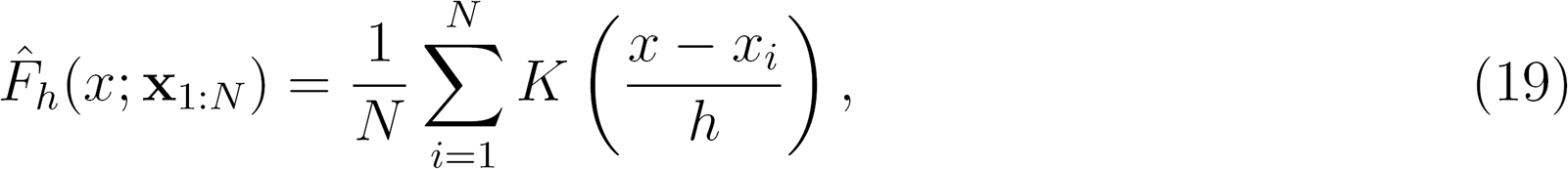

where 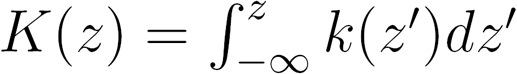 is an integrated kernel function and h is a bandwidth parameter (Simonoff, 1995, 1996; Simonoff & Tutz, 2000). We discuss later how this might be implemented via imperfect discrimination between items. From a coding perspective, smoothing can counteract the restricted resolution problem by spreading out stimulus representations to better cover the entire space, and can address the sampling variability problem by reducing the variance of the rank estimate. These correspond roughly to two canonical uses of smoothing in statistics: to produce estimates in regions where no data exists, and to improve estimates where data points are sparse (Burman, 1987).^5^

Figures 3 and 4 depict the effects of smoothing on the response distribution, and the entropies that result. Smoothing has two effects on *H*(*y*) and *H*(*y*|*x*). At low levels of smoothing, the sharp peaks of the output distribution due to limited coverage are smoothed, which has a beneficial effect on *H*(*y*) and an adverse effect on *H*(*y*|*x*). At high levels of smoothing, the distribution is smoothed completely so that *H*(*y*) and *H*(*y*|*x*) both converge to 0, which is detrimental for the former and favorable for the latter. The information-maximizing level of smoothing balances these forces and tends to produce an output distribution closer to uniform. Simulations shown in Figure 5 indicate that the information-maximizing bandwidth is decreasing in the sample size, and increasing in the variance of the attribute distribution. Thus while the redundancy tradeoff does not perfectly translate into the classical bias-variance tradeoff, they exhibit similar qualitative properties; in both cases, smoothing is most effective when the sample is small and when the underlying distribution is variable.

**Figure 3:**
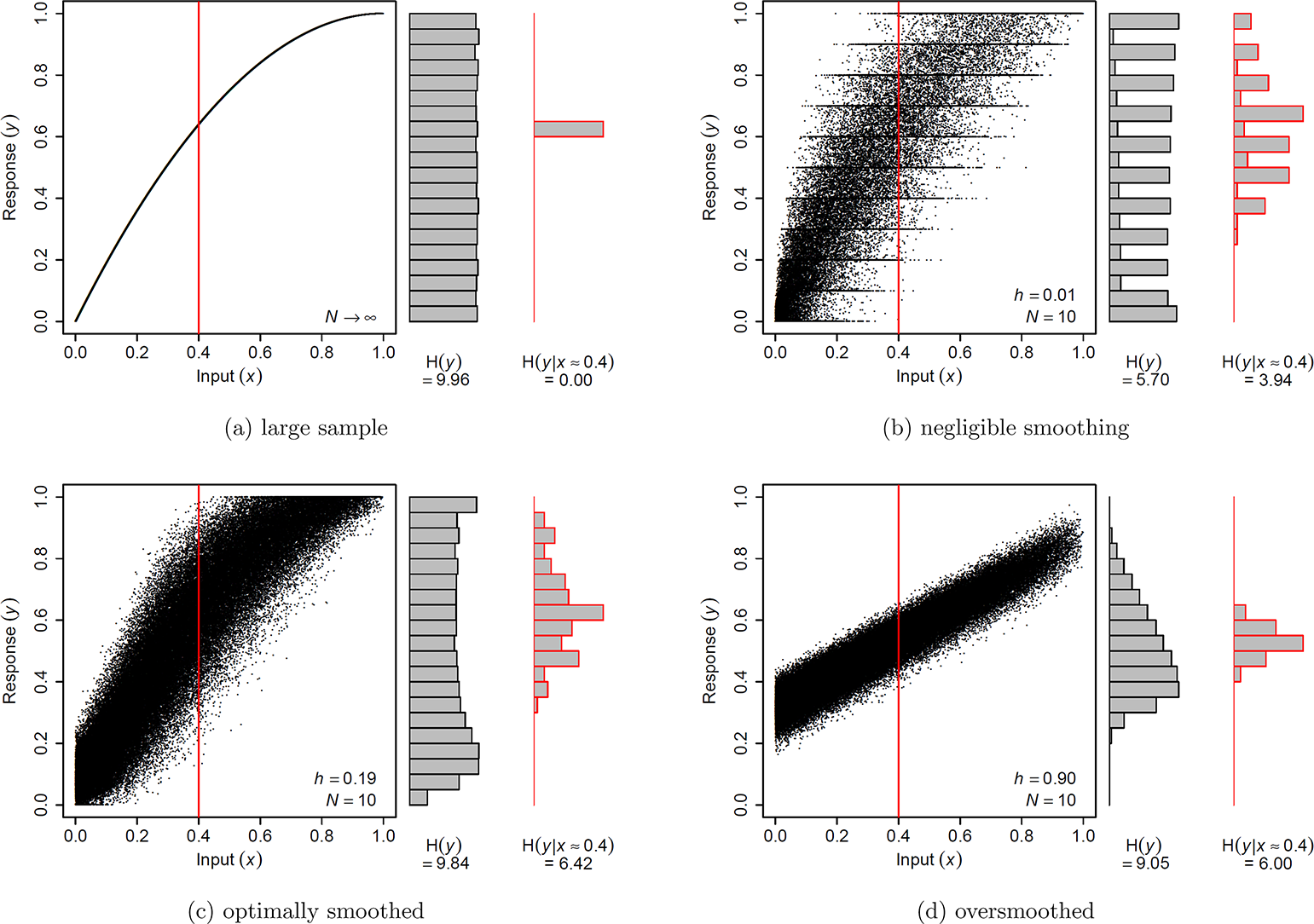
Effects of kernel smoothing on output entropy. Scatter plots depict samples from the joint distribution *f*(*x*,*y*), with 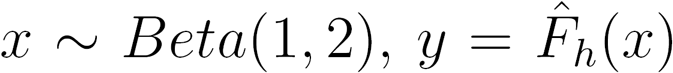, and a uniform kernel. Marginal histograms depict the distributions *f*(*y*) (black border) and *f*(*y*|*x* ≈ 0.4) (red border). **H*(*y*)* and *H(y\x)* are the entropy and conditional entropy in bits of the output *y*(*x* and *y* are discretized with bin width 1/1000).

**Figure 4:**
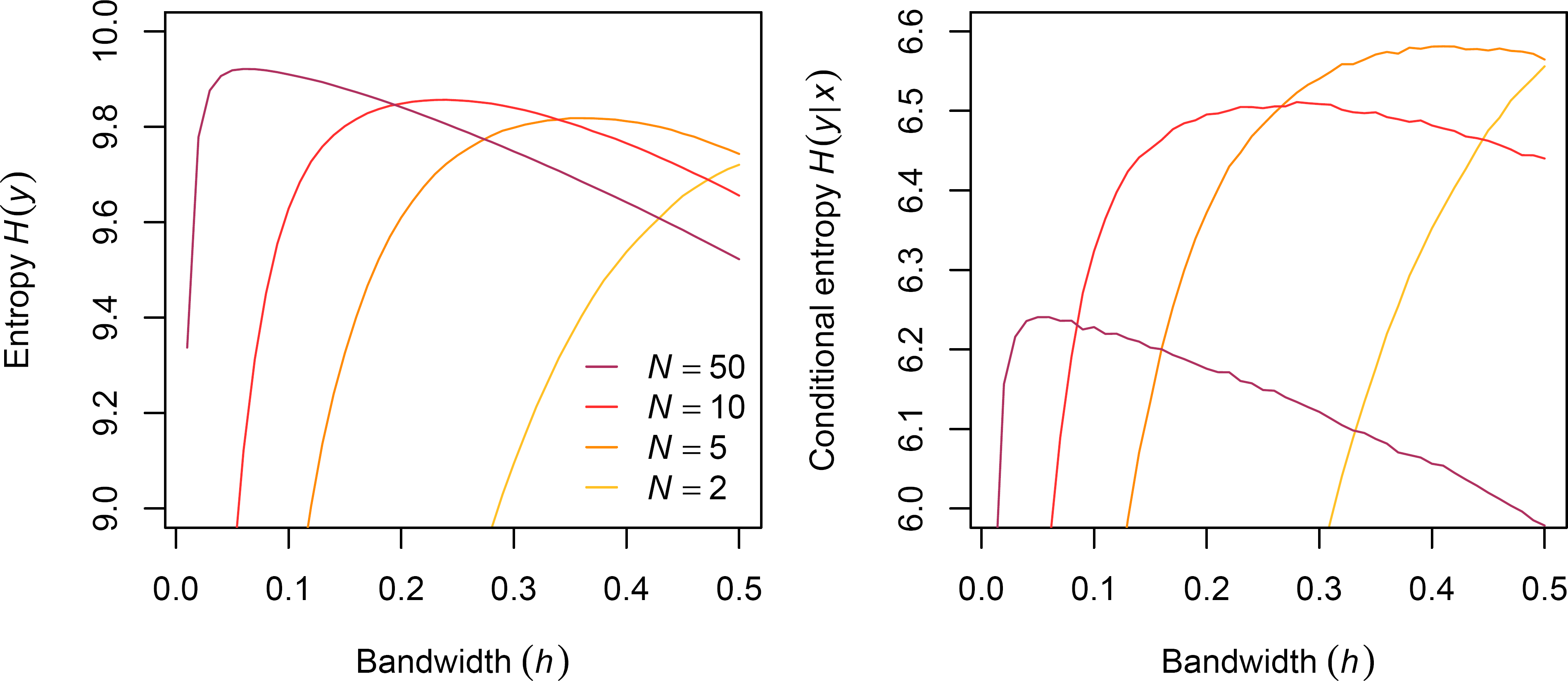
Smoothing increases entropy and conditional entropy at low levels and decreases them at high levels. Lines represent entropy and conditional entropy in bits (based on discretizations of *x* and *y* with bin width 1/1000). Attribute values *x* are drawn from a *Beta*(1, 2) distribution.

This analysis suggests that the principle of smoothing may guide the development and assessment of psychoeconomic models. In the following sections, we show that this leads to new insight into generalizations of DbS along with Parducci’s (1965; 1995) range-frequency theory.

## Range sensitivity as kernel smoothing

One limitation of DbS, noted by Stewart et al. (2006), is that it does not capture the effect of attribute range on decisions and other economic judgments. For example, increasing the range of gains causes people to become more risk-seeking (Lim, 1995), and to be less satisfied with a given gain (Parducci, 1968). Range effects have also been documented in wage satisfaction judgments (Brown et al., 2008; Tripp & Brown, 2016), university satisfaction ratings (Brown et al., 2015), lottery pricing (Blavatskyy & Köhler, 2009), and job application decisions (Highhouse, Luong, & Sarkar-Barney, 1999; Rynes, Schwab, & Heneman, 1983).

**Figure 5:**
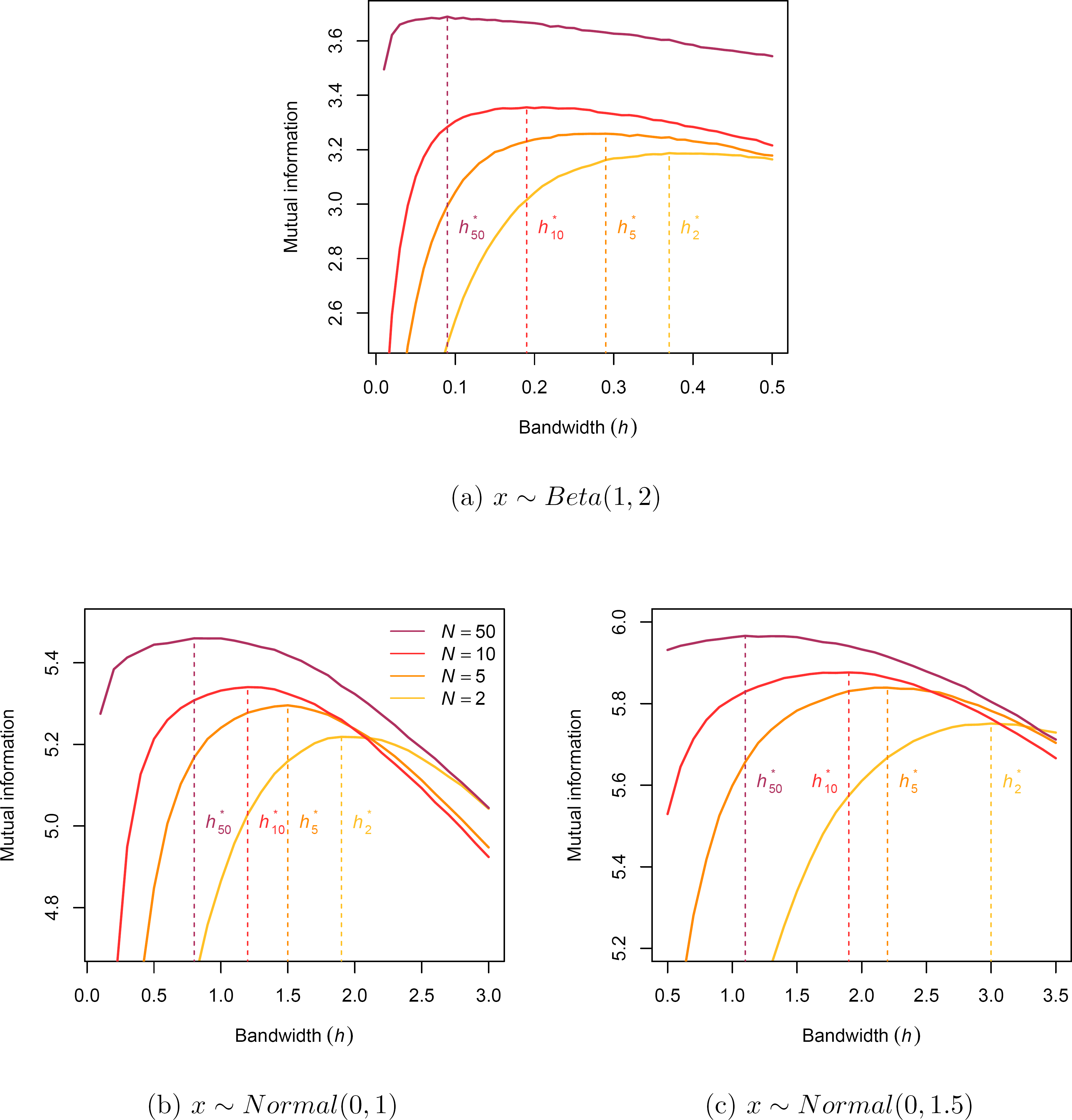
Optimal bandwidth *h\** is decreasing in sample size *N*, and increasing in variance of *x*. Lines represent mutual information in bits between *x* and *y* (both discretized with bin width 1/1000) as a function of uniform kernel bandwidth *h.*

Parducci’s (1995) range-frequency theory (RFT) is the most well-known account of these effects, grounded in the psychophysics of perception. The theory states that the judged magnitude (*J*) of an attribute value *x* is a convex combination of its sample rank (*F*, what Parducci refers to as its “frequency”) and its position relative to the range of attribute values (*R* = (*x* — *x_l_*)/(*x_h_* — *x_l_*), where *x_h_* and *x_l_* are the highest and lowest values):

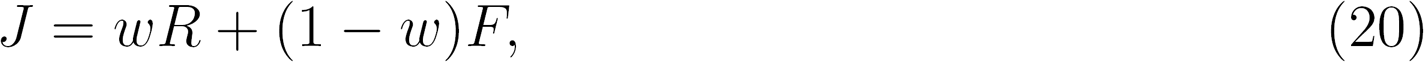

where ω is a weighting parameter that determines the compromise between range and frequency.

Parducci (1995) noted that the frequency component of RFT maximized information transmission, and informally suggested that the range component had error-abating properties. In line with this idea, we show that the range component of RFT can be derived as a kernel-smoothed estimate of the CDF using a uniform kernel. Both components are special cases based on the kernel’s bandwidth, and the weighting parameter heuristically tunes the degree of smoothing. The frequency component emerges when the bandwidth is 0, and the range component emerges when the bandwidth is proportional to the sample range. We accordingly propose that while the frequency component of range-frequency theory can be viewed as implementing redundancy minimization, the range component can be viewed as assisting with efficient coding in the face of noise.

Just as kernel density estimates can be written as the average of kernel densities *k*(*z*) centered around each data point, the distribution estimate can be similarly written as the average of the corresponding distributions 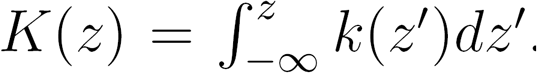. For a uniform kernel *k*(*z*) = 1/2 for |z| ≤1 and 0 otherwise, the distribution is *K*(*z*) = (*z* + 1)/2 for |*z*| ≤1, 0 for *z* <—1, and 1 for *z* > 1. Thus the CDF can be estimated smoothly by

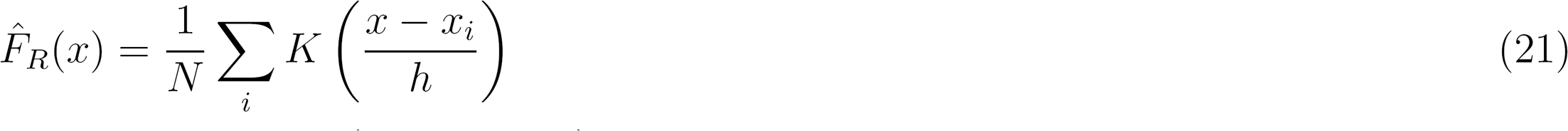

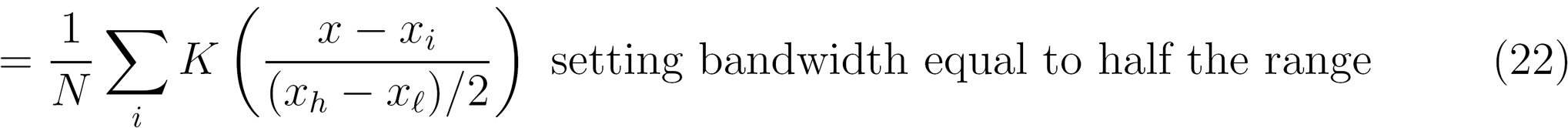

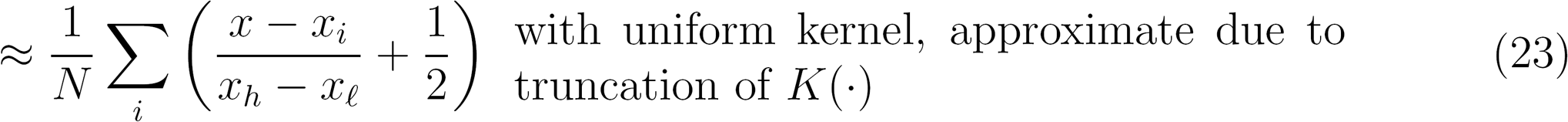

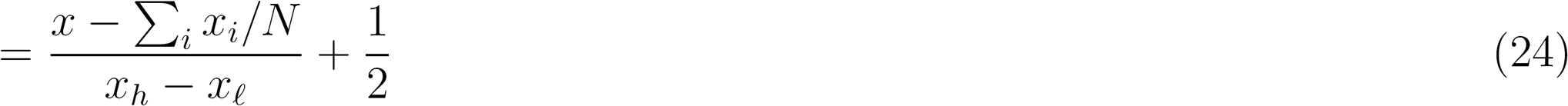

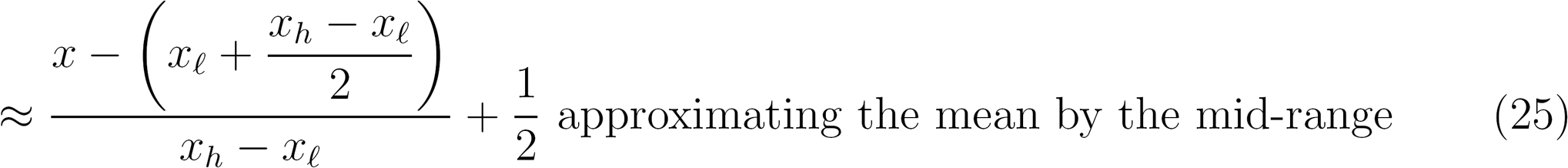

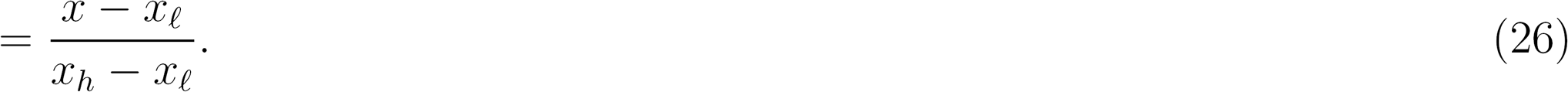

Hence, 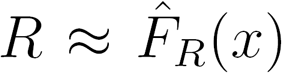, meaning the range component of RFT inherits properties of a kernel-smoothed estimate of the CDF.

Observe also that as bandwidth goes to 0, the estimate simply counts the number of data points less than the target value. If *x_i_* <*x* then 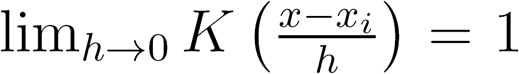, while if *x_i_* > *x* then 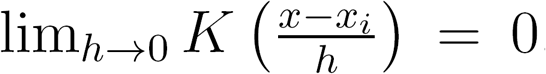. (In the knife-edge case that *x* = *x_i_*, 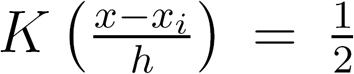.) This means the estimate becomes the unsmoothed empirical rank, so both the range and frequency components of RFT can be characterized as special cases of the kernel-smoothed estimate based on bandwidth. Thus the RFT prediction can be written equivalently as a combination of smoothed and unsmoothed distribution estimates:

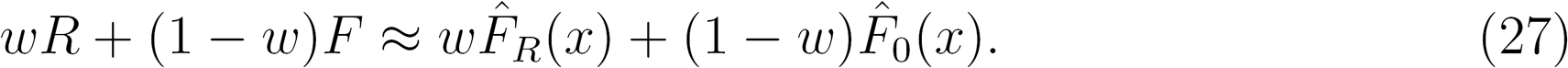

This derivation implies that the weighting parameter regulates the level of smoothing in place of the bandwidth parameter. If psychoeconomic functions are indeed attuned to the statistical properties of encountered attributes, the argument predicts that the range component should come to dominate as noise increases.

Why assume that the kernel is uniform and that bandwidth is proportional to range? These turn out to have optimal statistical properties for estimating CDFs. The uniform kernel minimizes mean integrated squared error when estimating distribution functions (Jones, 1990). Although other kernels are nearly as efficient, clearly the uniform is among those that work quite well. Optimal bandwidths for distribution estimation are typically related to standard deviation. This occurs because when standard deviation rises, data becomes sparse and variance in the estimate increases. The benefit of variance reduction rises relative to the cost of increased bias from a higher bandwidth. Distribution-optimal plug-in bandwidths generally take the form 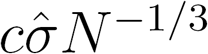 (Azzalini, 1981; Hansen, 2004) for various constants *c*. Standard deviation is approximately proportional to range, as illustrated by the “range rule” which states that the standard deviation is roughly the range divided by 4.

The plug-in bandwidth also directly reveals that the sample size should influence bandwidth. Smoothing is only useful when samples are small, otherwise the bias it induces outweighs the reduction in variance, and increases error on net. This argument suggests that the weighting parameter which stands in for the bandwidth should also vary with the relevant sample size (whether of the observed context or the subsample drawn from memory). Although the mathematical expressions we present are not strictly optimal in an information-theoretic sense, they capture some important regularities borne out by simulation, namely that smoothing should increase when the variability of the attribute values is large and should decrease with the number of values retrieved from memory. Moreover, the optimal level of smoothing may change according to other factors that are less obvious, such as the granularity of response categories, which we illustrate in the following section.

## Predicting range sensitivity

In the above characterization of RFT as a smoothed distribution estimator, the range weighting parameter w controls the degree of smoothing. This parameter is typically fixed at 0.5 when applying RFT to the data. However, the smoothing interpretation generates new predictions about what should cause it to vary. Here is one example of how this can help account for observed data.

When stimuli or responses are limited to a finite number of categories, people tend to partition the space so as to evenly use all bins (in other words, their limens are the distribution quantiles). This is just as redundancy minimization demands. Uniform use of categories has been found in judgments of loudness (Stevens, 1958), size (Parducci & Perrett, 1971), and could also explain the “numbers-of-levels” effect in marketing, whereby an increase in the number of attribute levels leads to an increase in the relative importance of that attribute (Verlegh, Schifferstein, & Wittink, 2002). However, Parducci and Wedell (1986) further find that the number of available response categories influences the effects of skewness and the apparent weighting of range and frequency components. As the number of categories increases, the range component becomes more dominant and skewness has a diminished effect on judgment.

Optimal smoothing provides a possible explanation for this result. We have seen the damage caused by finite samples when trying to uniformly distribute stimuli across many representational states (equivalently, categories); the code cannot finely partition the space, and the representations are quite noisy. However, these problems are less severe when few response categories are available, because coarsening the response space naturally mitigates the effects of noise (Simonoff & Tutz, 2000). For example, with *N* = 1, the quantiles needed to partition a high resolution space are poorly estimated; in contrast, the two-category partition constitutes a median split, and the lone draw is a passable estimate of the median. In the regime where *M* < *N* (with simulations depicted in Figure 6), the response resolution, and hence *H*(*y*), is already capped based on the categories rather than the sample. Moreover, the channel noise *H*(*y*|*x*) starts decreasing in sample size because the resolution remains fixed while the quantiles are estimated more precisely. These finite sample issues are thus less troublesome with fewer categories, reducing the possible upside of smoothing.

**Figure 6:**
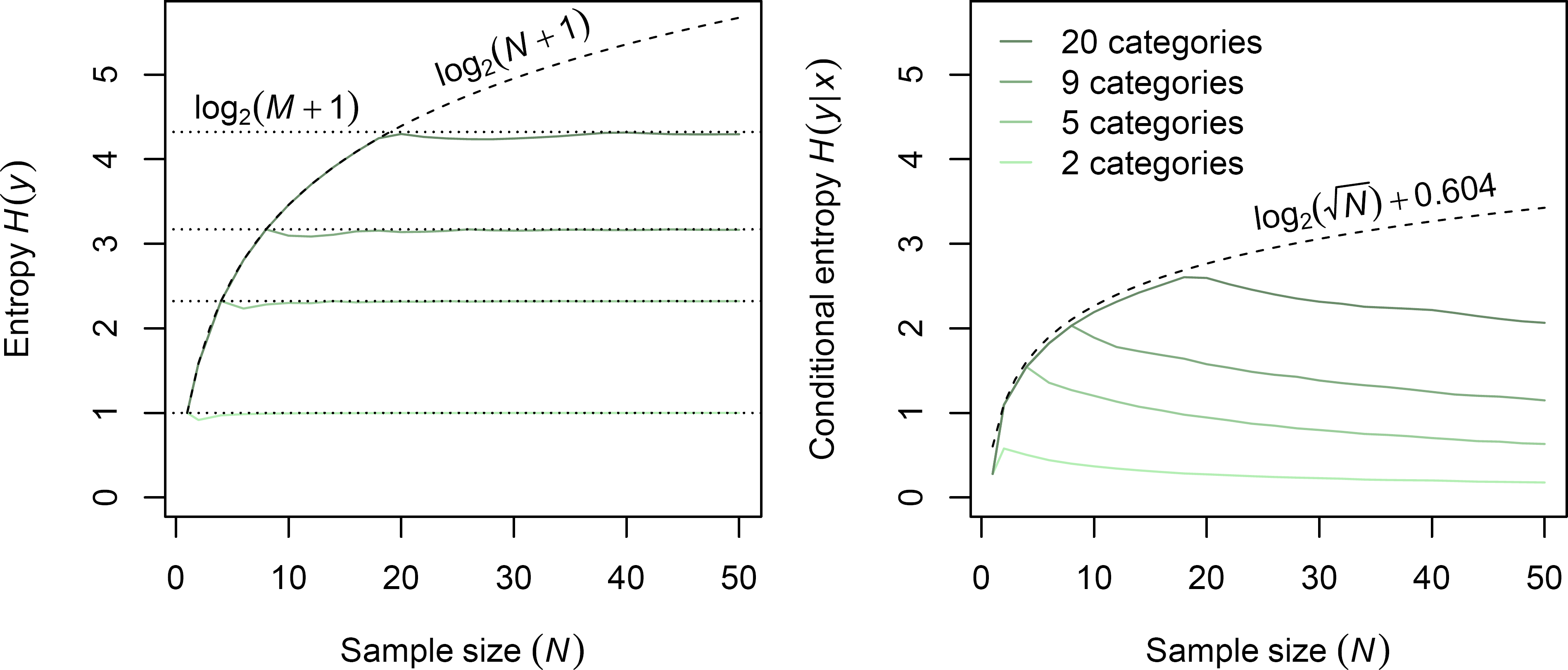
When the number of categories is small, the adverse effects of finite samples are naturally abated. Lines represent entropy and conditional entropy in bits, based on *y* and a discretization of *x* with bin width yiooo. Attribute values *x* follow a *Beta(1,* 2) distribution.

To illustrate the predictions resulting from the smoothing interpretation of RFT, we conduct simulations encoding *y* based on an RFT coding scheme, and show how mutual information varies with the range weight *w*, the number of response categories, and the sample size. Following Parducci (1965), RFT limens are the weighted average of range and frequency limens. Range limens here are the quantiles of the standard uniform distribution, and frequency limens are the linearly interpolated quantiles of the sample (made to include the endpoints 0 and 1). Which quantiles are chosen depends on the number of response categories. With three categories, for example, an even partition is formed by the 33rd and 67th percentiles. We pick as the context a skewed Beta distribution with parameters *α* = 1 and *β*= 2 (or equivalently, vice versa) to roughly resemble Parducci and Wedell’s (1986) stimuli.^6^

**Figure 7:**
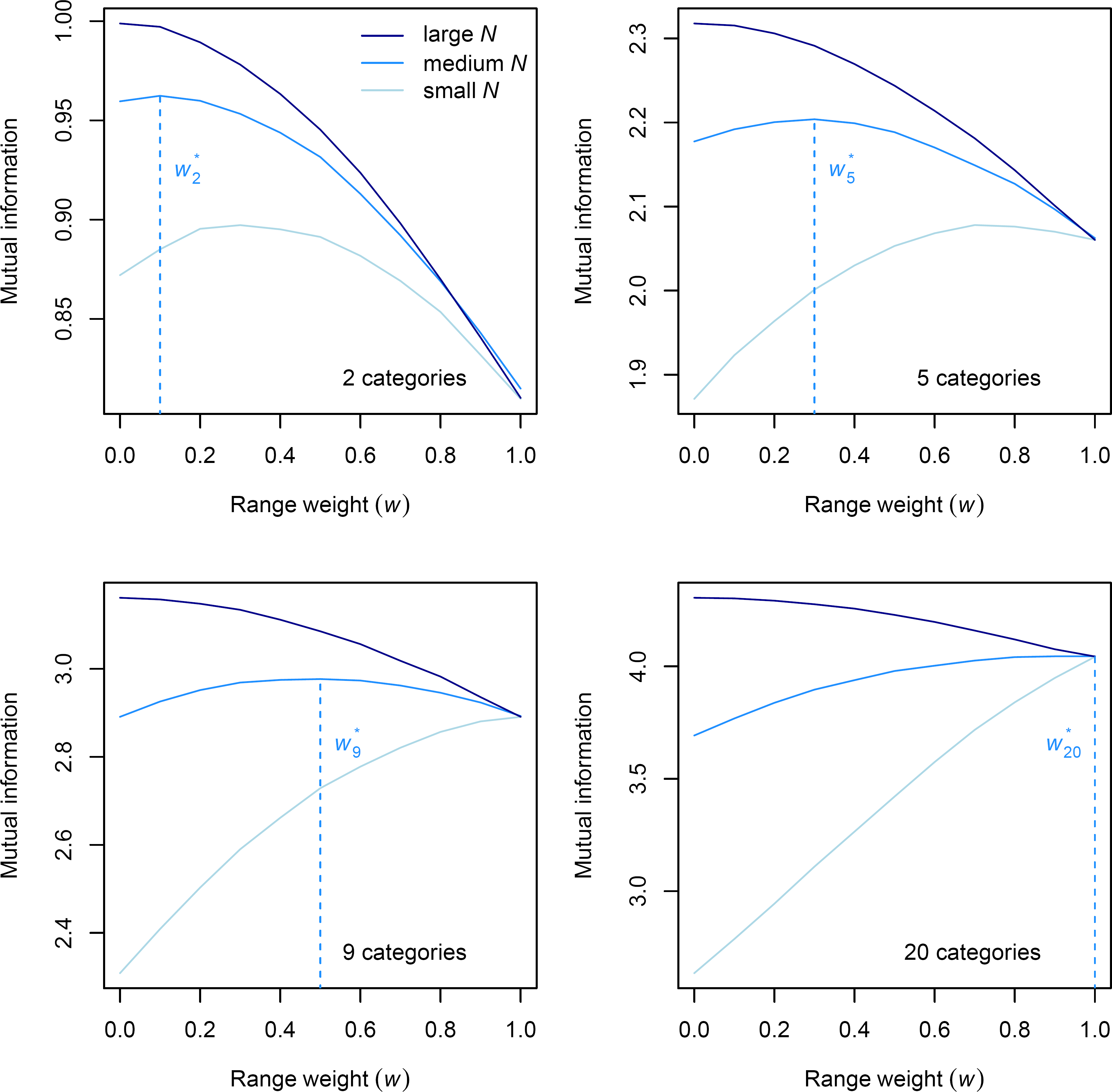
Optimal range-frequency theory weight w* is increasing in number of response categories, and decreasing in sample size. Lines represent mutual information in bits between *x* and *y* from RFT (x discretized with bin width 1/1000) as a function of range weight parameter *w*. Attribute values *x* follow a *Beta*(1, 2) distribution.

Figure 7 indicates that the information-maximizing range weight is increasing in the number of categories, as predicted. Similar to the earlier simulations, the optimal range weight is decreasing in the sample size, further supporting the idea that it acts as a smoothing parameter. The utility of smoothing is a consequence of finite samples, so with a large sample, the optimal weights shrink substantially and the range component becomes negligible however many categories are available. Note that in order to display the results clearly, the sample sizes are taken to be *N* = 10^2^, 10^3^, and 10^5^. These values are greater than would be expected if they are meant to reflect the number of items retrieved from memory. However, the sample sizes required for interior solutions (i.e., w* ≠ 0,1) depend on the exact distributions and functional forms assumed, and are in some cases more psychologically plausible. More importantly, the same qualitative results hold across various sample sizes, category numbers, and distribution parameters, demonstrating that the information-enhancing properties of the range component vary as predicted by principles of smoothing.

## Smoothing as reduced discriminability

As suggested by Stewart et al. (2006), and later elaborated by Brown and Matthews (2011), range-like effects can be captured by DbS if one assumes that experienced attribute values are not perfectly discriminable in memory. In particular, many models of memory assume that the discriminability of items is inversely proportional to their density in attribute space (e.g., Brown, Neath, & Chater, 2007). Thus, an item is less likely to be retrieved if other similar items enter into competition for retrieval. This mechanically flattens out the effective retrieval distribution, damping its original skewness. Estimated ranks are then based on a more uniform distribution, mimicking an increase in the relative importance of the RFT range component. Parducci and Wedell (1986) find they can also explain their category results described in the previous section using a model that acts like that of Brown and Matthews. Reducing stimulus discriminability and increasing range weight thus have similar explanatory capabilities.

This link might explain why range weighting appears to change depending on the salience of the contextual distribution. For example, when the distribution must be drawn from memory due to sequential rather than simultaneous presentation, the range component seems to become more important (Choplin & Wedell, 2014; Niedrich, Sharma, & Wedell, 2001; Qian & Brown, 2005). Recalling samples from memory injects noise and reduces discriminability. Similarly, customers who have been exposed to a trend in prices exhibit less range weighting (Niedrich, Weathers, Hill, & Bell, 2009), perhaps because the clearer structure of the contextual distribution enables more precise retrieval.

The efficient coding framework offers another perspective on this connection: imperfect discrimination may be a mechanism for introducing redundancy to reduce coding errors. Kernel smoothing from a sampling perspective can present as reduced discriminability.

Suppose that when an item is drawn from memory, some uncertainty is felt about its true location. This entails that items won’t be completely distinguishable, and those nearer each other will be harder to distinguish. These are the assumptions imposed by reduced discriminability models. The coarse binary comparisons of DbS are then replaced with graded assessments of order to allow some tolerance. Rather than simply determining whether the target is greater than each sample value, the differences between the target and the samples are judged as significant to varying degrees based on the level of uncertainty.

This smoothed comparison is exactly what a kernel encodes, illustrated in Figure 8. Each sample value *x_i_* contributes 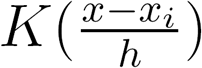 to the rank estimate of *x*. The level of uncertainty is represented by the bandwidth *h*, and accordingly controls the degree of smoothing. If this is small, the contribution of *x_i_* boils down to 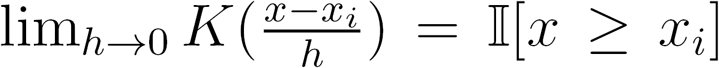, a pure binary comparison. As uncertainty grows large, this becomes 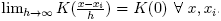. No comparisons can be made at all, so the effective distribution becomes more uniform, just as in the reduced discriminability models. The optimal amount of smoothing to account for sampling variability is somewhere in the middle, informed by range as discussed earlier. In sampling terms, range provides information about how much comparison tolerance should be allotted. With a discrete response space, the comparisons exhibit some granularity, but the essential smoothing properties are shared by discrete analogues of the continuous kernel (Simonoff, 1995, 1996).

**Figure 8:**
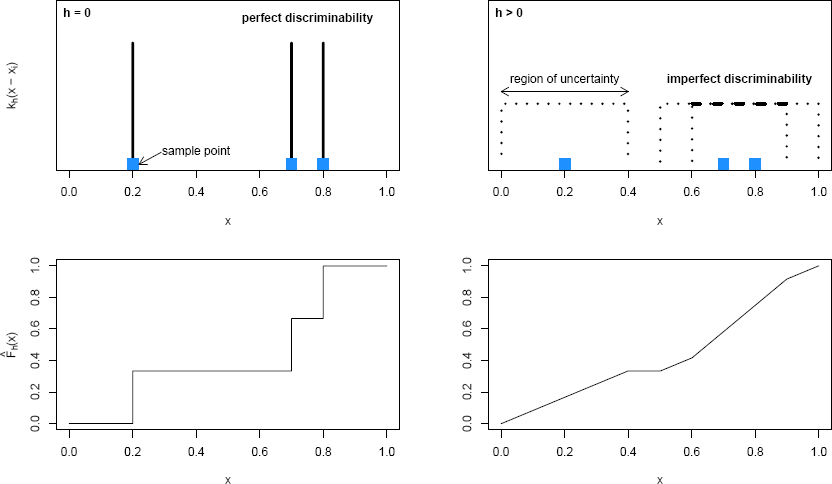
Kernel smoothing presents as imperfect discriminability. (left) When items are completely distinguished from each other, bandwidth is zero, and no distribution smoothing occurs, (right) When some tolerance is allowed for uncertainty in location, bandwidth is positive, and the distribution is smoothed.

We note further that the region of uncertainty can itself be instantiated by resampling each retrieved item, suggesting how range sensitivity could be implemented via purely binary comparisons. If uncertainty is high, resamples will be spread out, some of which will be greater than items otherwise higher on the scale. This may be more cost-effective than drawing fresh samples if resamples are cheaper to obtain, which is plausible when they can be anchored to their originally retrieved estimates. As the cognitive simplicity of DbS is a key part of its motivation, placing extensions on equal footing contributes to their justification. In addition, the range component of RFT requires recall of only the distribution endpoints, and thus may be cognitively undemanding in that sense. Kernel-smoothed rank need not be more taxing to compute than empirical rank.

Thus the assumption of imperfect discriminability can be directly tied to the implementation of kernel smoothing. From the efficient coding perspective, the relationship between range sensitivity and imperfect discriminability is not so surprising. They are alternative facets of the same smoothing phenomenon.

## Smoothing and context effects

The explanatory power of DbS stems from its ability to predict how judgment is influenced by the contextual distribution. Context distorts value judgments in ways that produce violations of classical expected utility theory such as those described earlier. Incorporating smoothing enables DbS to capture an even wider class of context effects. We have discussed some of these in a single-attribute setting in terms of range sensitivity.

In a multi-attribute setting, Ronayne and Brown (2017) show how DbS with a form of local sampling can predict the attraction, compromise, and similarity effects. The compromise effect is most closely linked to locality so we focus on this. It refers to a scenario in which an extreme option makes the intermediate option more likely to be chosen. Figure 9 illustrates a kernel smoothing version of Ronayne and Brown’s setup. Options *A* and *B* are focal price-quality pairs, and *R_A_* and *R_B_* denote the regions they uniquely dominate. *C_A_* denotes the third, extreme option that makes *A* the compromise and hence preferred. The model supposes that a finite sample is drawn from memory, and the subjective value of each option is the number of samples in the region that it dominates. Hence their unique dominance regions play a key role in the relative values of options.

Crucially, Ronayne and Brown assume that regions close to the presented options in attribute space are preferentially sampled; the support of the effective retrieval distribution is represented by the shaded region. As a result, *C_A_* contributes more probability mass to the solo-dominance region of its neighbor *A* than it does to that of *B*. Thus the presence of *C_A_* benefits *A* relative to *B*, producing the compromise effect.

This locality assumption can be interpreted as a form of kernel smoothing. If the available options constitute draws from the context, the location they represent for purposes of rank comparison is inexact. Along similar lines to the previous section, the resulting region of uncertainty instantiates a kernel which trails off when it gets farther from the option. If this is the case, optimal smoothing predicts that the size of the kernel should grow according to the scale of the option distribution, naturally providing a reason for compromise effects at all scales. The scope of the compromise effect should also be related to factors like the number of options available and the psychological noisiness of context retrieval. This could help explain why the compromise effect decreases when more options are presented (Gourville & Soman, 2007). Because a larger sample is available, smoothing is less necessary and bandwidth decreases.

**Figure 9:**
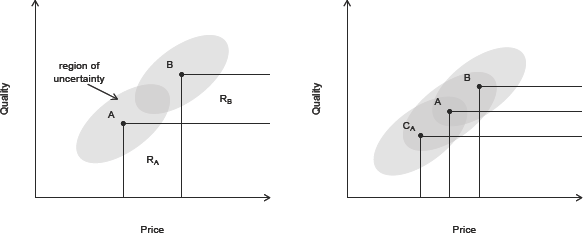
Compromise effects can be generated by multi-attribute DbS with locality. (left) *A* and *B* are the focal options, and *R_A_* and *R_B_* represent their unique dominance regions. Locality of sampling can be interpreted as a kernel around each option. (right) The third option *C_A_* contributes more mass to *R_A_* than *R_B_*, benefiting *A* over *B*.

## Neural evidence and implementation

There is much evidence of neuronal gain control that produces range sensitivity (e.g., Padoa-Schioppa, 2009; Tremblay & Schultz, 1999), though little research has attempted to pinpoint value encoding by option rank. Hence this remains an important area for further study.

Mullett and Tunney (2013) provide the most direct neural evidence to date. In their experiment, participants were faced with blocks containing either rewards of £0.10, £0.20, and £0.30 or rewards of £5.00, £7.00, and £10.00. Using fMRI, they found activity in the ventromedial prefrontal cortex and anterior cingulate cortex to be linear in the global option rank rather than absolute value, meaning the difference between £5.00 and £0.30 was similar to the difference between £0.30 and £0.20. Interestingly, these regions encoded rank based on the set of stimuli presented across the whole experiment, while activity in the caudate and thalamus scaled according to the experimental block, exhibiting a more local context dependency.

Why have there not been more direct signs of rank encoding? We note two possible reasons why rank-based value representations may be difficult to discern.

First, the point of smoothing is to gracefully approximate rank across the entire range of values. In general, this mechanism reduces sensitivity to contextual skew, as illustrated by RFT, according to which the response function is a convex combination of the empirical CDF (frequency component) and a linear function (range component). Taken to the pure range extreme, this leads to linearity across the support of the distribution. Thus, smoothing could provide a normative rationale (consistent with efficient coding) for why Rustichini, Conen, Cai, and Padoa-Schioppa (2017) observe that offer value cells in the orbitofrontal cortex adapt to stimulus range, and yet exhibit quasi-linear tuning curves even when the value distribution is non-uniform.

Second, rank-based coding may take unusual forms not previously considered by those studying decision making. In practice, forms of rate and population coding have dominated the neuroeconomic literature. Alternatively, efficient coding could be naturally implemented in the temporal domain. On a neuronal level, encoding based on the rank of spike timing— known as rank order coding (Thorpe & Gautrais, 1998)—has several benefits. Such schemes convey information more efficiently than standard rate codes while using simpler and more robust decoding mechanisms than precise timing codes (VanRullen & Thorpe, 2001). The most important information can be transmitted first and quickly decoded, enabling the kind of swift responses to stimuli that may be essential for survival. Rank order coding is capable of transmitting information on the rapid timescales of the sensory domain (VanRullen, Guyonneau, & Thorpe, 2005), and there is evidence of its role in retinal processing (Portelli et al., 2016).

We show how smoothing can be naturally integrated into rank order coding. Existing computational implementations permit neurally plausible extensions that would generate smoothing. Consider the simple example circuit shown in Figure 10 modeled after Thorpe, Delorme, and Van Rullen (2001). Suppose neurons *A*, *B*, and *C* represent stimuli with attribute intensities *A > B > C*. They are to be compared using their rank-influenced cumulative outputs *a,* *b*, and *c*. Normally *A* will fire first because the intensity of its stimulation is highest, *B* will fire later, and *C* will fire later yet. So a should be the greatest, and this can be accomplished straightforwardly via the inhibitory interneurons *I* that attenuate the effects of later firing inputs. Because *A* fires first, the inhibition will grow to depress the effects of *B*, and subsequently *C*, reflecting their lower rank. Stronger inhibitory power sharply raises the relative value of the highest ranks, producing a value of a much greater than b and c.

In this mechanism, neural activity is implicitly assumed to decay instantly. However, a gradual decay may be more realistic. If active traces remain after a neuron’s initial spike, subsequent neurons can fire before earlier traces completely decay. The inhibitory effects of higher ranked neurons are accordingly lessened since part of their activity occurs after lower ranked neurons spike. This means the active trace operates like an asymmetric kernel, with the decay distribution affecting the kernel shape and the decay rate controlling the bandwidth. A slow decay reflects a large bandwidth that smooths the encoded attribute ranks. Steep changes due to differences in rank are consequently diminished.

In summary, we have shown how efficient coding could be implemented naturally in the temporal domain, with smoothing implemented through gradual decay of activity. However, this hypothesis remains to be verified experimentally.

**Figure 10:**
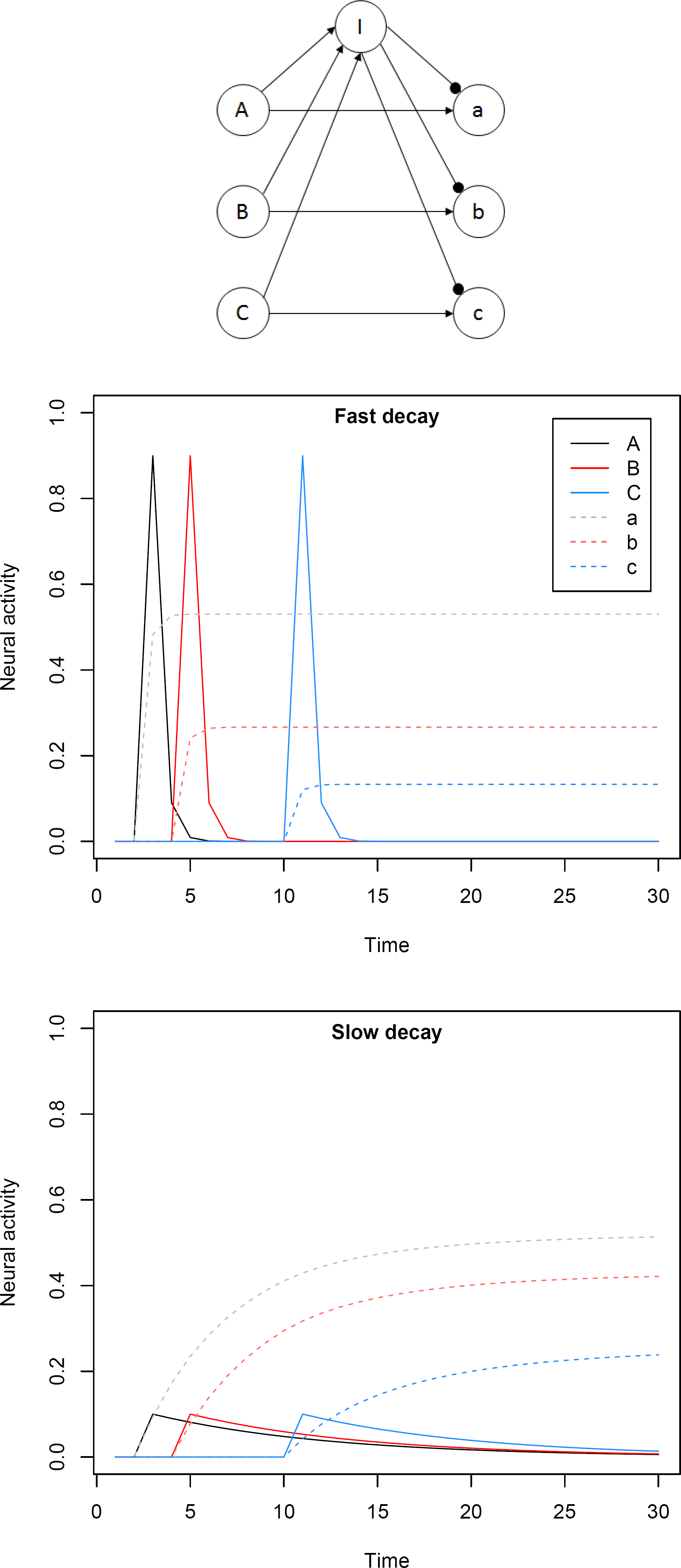
Slow post-spike activity decay implements smoothed rank order coding. (top) Diagram of simple network implementing rank order coding. Simulation assumes divisive inhibition which attenuates effect of inputs by a factor of 0.5^*I*/(*t*)^. (middle) When decay is fast, output values are sharply sensitive to rank, as *a >> b, c.* (bottom) When decay is slow, output values are less sharply sensitive to rank, as the decay operates similar to an asymmetric kernel.

## Neuroeconomic models and efficient coding

We provide an adaptive rationale for DbS and its family by grounding them in the efficient coding framework. This elevates DbS to a select class of theories that have been justified both normatively based on principles of efficient coding and descriptively based on data from perceptual as well as economic domains. It is no coincidence that there is a link between these elements. From the start, research in behavioral economics was guided by the notion that cognitive processes underlying judgment resembled those underlying perception (Kahneman, 2002). This supposition works well partly because efficient coding constitutes a unifying principle that spans low-level perception and high-level judgment.

Natural selection imposes pressure on organisms to efficiently represent the information they need to survive and reproduce. The brain spends a tremendous amount of energy, accounting for 20% of resting oxygen consumption in adult humans, most of which is directly required for signaling (Laughlin, 2001). Given the enormous metabolic costs of neural activity and infrastructure, encoding information wastefully would produce a steep drop in fitness, and should therefore be sharply curtailed by selection pressures. This argument applies generally across species and types of processing.

The realm of perception provides a low-level testbed in which computational descriptions of problems to be solved, and hence the nature of optimality within them, can be more transparently specified. Thus the development and assessment of theories, including those based on efficient coding, can progress at a faster rate. This creates an arbitrage opportunity for the study of decision making. Research in cognitive science and computational neuroscience has yielded a wealth of precise, quantitative, and tractable characterizations of perceptual processing. These insights are being ported into the study of decision making and have already demonstrated great predictive power.

To better locate DbS in this landscape, we describe two other branches of neuroeconomic modeling which are motivated by their optimal information-theoretic properties under alternative assumptions.

First, our analysis ignored statistical dependencies between the symbols that comprise the representation (e.g., the ones and zeros of the codes in Figure 1), making redundancy purely a matter of unequal symbol usage (sometimes called first-order redundancy). This would be appropriate if, for example, the representation were implemented by a single neuron with many states, or by many neurons firing one at a time. However, if multiple neurons were collectively responsible for encoding a complex stimulus, then higher-order redundancy could arise from interdependencies between their individual representations (Atick, 1992; Brady & Field, 2000). This issue occurs often in visual processing, as neighboring pixels in natural scenes exhibit highly correlated luminance values, producing higher-order redundancy in pixel-based codes. One way to address this problem is by local suppression of activity, and this is the approach taken by divisive normalization.

Divisive normalization is a mechanism whereby neighboring neurons inhibit each other and thus exhibit responses that are normalized with respect to their pooled inputs. This leads to adaptive gain control that calibrates the sensitivity of neuronal responses according to the local context. It was proposed to assist with redundancy reduction in sensory processing (Schwartz & Simoncelli, 2001a; Wainwright, Schwartz, & Simoncelli, 2002), and has been shown theoretically and empirically to help reduce higher-order redundancies in such settings. For example, it removes statistical dependencies in representations of the heavy-tailed multivariate distributions that characterize natural image statistics (Lyu, 2010, 2011; Malo & Laparra, 2010). Divisive normalization has been observed across various species in neural pathways for vision (Busse, Wade, & Carandini, 2009; Carandini, Heeger, & Movshon, 1997; Heeger, 1992), audition (Rabinowitz, Willmore, Schnupp, & King, 2011; Schwartz & Simoncelli, 2001b), olfaction (Luo, Axel, & Abbott, 2010; Olsen, Bhandawat, & Wilson, 2010), and even multisensory integration (Ohshiro, Angelaki, & DeAngelis, 2011). It is considered a “canonical computation” due to its prevalence (Carandini & Heeger, 2012). When applied to economic behavior, divisive normalization predicts context dependence that is able to account for classic deviations from expected utility theory (Louie et al., 2011, 2013, 2014; Rangel & Clithero, 2012).

A second assumption in our analysis was that the organism could not reduce the cost of information transmission by strategically changing the signal-to-noise ratio. This is reasonable when precision is of vital importance. However, giving up the ability to crisply distinguish between certain values can be adaptive at times. If this were feasible, it could be worthwhile for the organism to transmit, say, only the first bit of the codes in Figure 1 when the cost of full precision outstrips the benefits (the calculation of which could involve consideration of downstream processing or behavioral responses). Sequential sampling models elaborate on this approach with a noisy channel, and describe the process of decision making as the optimal accumulation of evidence over time pertaining to different options (Bogacz, Brown, Moehlis, Holmes, & Cohen, 2006; Laming, 1968; Stone, 1960).

In typical sequential sampling models, an agent is trying to identify the unknown state of the world based on noisy signals received over time, and an option is chosen when the evidence accumulated in its favor reaches some threshold. This threshold was originally derived from efficient Bayes-optimal statistical algorithms for estimating the state of the world from a sequence of data generated at a unit cost (Arrow, Blackwell, & Girshick, 1949; Fudenberg, Strack, & Strzalecki, 2017; Wald, 1947; Wald & Wolfowitz, 1948). These models closely match the joint distributions of choices and response times observed in perceptual tasks (e.g., Ratcliff, 1978, 2002; Ratcliff & Rouder, 1998; Smith, Ratcliff, & Wolfgang, 2004), as well as patterns of neural activity (Gold & Shadlen, 2002; Hanes & Schall, 1996; Ratcliff, Cherian, & Segraves, 2003; Shadlen & Newsome, 2001; Smith & Ratcliff, 2004). Woodford (2014, 2016) provides a complementary information-theoretic formulation in which the threshold rule is assumed, and instead the degree of noise in the code can be controlled at a cost proportional to information-processing capacity. He derives the dynamic signal precision which maximizes expected utility (based on the reward for correct identification), finding it to depend (negatively) on the experienced evidence gap between options. Woodford’s theory applied to economic decisions predicts aspects of stochastic choice even more accurately than other sequential sampling models which assume a fixed-rate accumulation process driven by the value difference between options (Krajbich, Armel, & Rangel, 2010; Krajbich, Hare, Bartling, Morishima, & Fehr, 2015; Krajbich, Lu, Camerer, & Rangel, 2012; Krajbich & Rangel, 2011; Milosavljevic, Malmaud, Huth, Koch, & Rangel, 2010).

Thus, efficient coding provides a unified set of principles for understanding many kinds of cognitive processes including economic decision making. It helps account for apparent violations of rational choice by recourse to deeper forms of optimality that reflect the cost-effective distortion of mental representations. Given its deep foundations and broad applicability, this approach will doubtless continue to be a fruitful source of neuroeconomic theories in the future.

## Conclusion

Abundant evidence demonstrates context sensitivity in judgment and decision making that deviates from classic theoretical models. Decision by sampling was intended to account for such data, proposing that the value of an attribute is encoded as its rank in a contextual distribution drawn from memory. This can be computed by tallying ordinal comparisons between the target attribute level and samples from the context, and entails that psychoeconomic functions are intrinsically malleable. Here, we ground DbS in an influential strand of theoretical neuroscience which posits that activity patterns in the brain efficiently represent information. This hypothesis of efficient coding predicts that neural processing, and therefore the psychophysical functions it generates, should be adapted to natural stimuli faced by the organism.

We identified DbS (and equivalently, the frequency component of range-frequency theory) as an implementation of information-theoretic redundancy minimization, which is efficient in a noiseless communication channel. Redundancy minimization requires values to be encoded so that they are uniformly distributed across the bounded response range, which is achieved by the rank transformation of DbS. However, when only a finite sample can be drawn from memory, as is made necessary by inherent computational constraints, the transmission of information is impaired. We theoretically specified the problems caused by finite samples, and showed that smoothing could help counteract these problems and partially restore coding efficiency.

We drew out the implications of this smoothing, showing that under certain assumptions that reflect optimal smoothing, the range component of RFT can be derived as a kernel-smoothed estimate of rank. This derivation revealed that RFT implements efficient coding in a previously unrecognized fashion. It also suggests that principles of optimal smoothing enable us to predict variation in the RFT weighting parameter, and we demonstrated that this can help account for past data on how judgment is affected by the number of available response categories. Psychologically, kernel smoothing can manifest as reduced discriminability of retrieved items, which sheds light on how previous extensions of DbS and RFT that assume imperfect discriminability capture range sensitivity. Similarly, extensions of DbS based on locality of sampling that capture context effects observed in decision making can be interpreted as kernel smoothing, providing further rationale for such approaches.

These insights into DbS open up many directions for future research. First, our analysis indicates that the optimal amount of smoothing depends on sample size; thus, can we predict smoothing from individual differences in memory capacity? Second, can we manipulate the degree of smoothing by changing parameters of the task? For example, can we increase smoothing by placing individuals under cognitive load? Third, can we find direct evidence for adaptive smoothing in the brain? Answering this question will require measurement techniques with high temporal resolution (such as single unit recordings) in order to test the predictions of the temporal coding scheme described in this paper. Finally, can we link this coding scheme to decision making behavior, such as context effects? We believe that efficient coding provides a powerful framework for addressing these questions.

## Footnotes

1 Although we separate *x* from the sample in our notation, this should not imply that the target itself is necessarily excluded from the sample.

2 Strictly speaking, this describes what is called higher-order redundancy, though our approach focuses on firrst-order redundancy. We will discuss this distinction in the penultimate section of the paper

3 We note two implicit assumptions in this analysis. First, the judged magnitude is equal to the neural response *y* rather than some further transformation (i.e., the decoding function is linear). Second, the optimization criterion is insensitive to costs other than expected codeword length. Relaxing these assumptions is beyond the scope of the present paper, which only proposes a theory of optimal encoding and not decoding, and shows that this is able to explain the shapes of psychoeconomic functions based on the shapes of the priors. However, different analyses could lead to further insights as demonstrated by Wei and Stocker (2015, 2017) and Park and Pillow (2017) in perception and Woodford (2012a, 2012b) in decision making.

4 While conditional entropy is increasing in *N* when *M* > *N*, the opposite occurs when *M* < *N*, as will become important later. Thus *H*(*y*|*x*) is able to converge to 0 when *N* → ∞ for a fixed *M*.

5 An alternative solution might be to estimate a parametric model of the attribute distribution using the sample (in contrast to our nonparametric solution). Brown, Wood, Ogden, and Maltby (2015) propose that this can help account for apparent range effects of the sort we discuss later. While this is an intriguing possibility, such an approach would require the agent to have access to a level of distributional knowledge that we have assumed is unavailable. Moreover, it would require specification of how the distributional model varies across contexts, adding degrees of freedom.

6 The results still hold with other parameter values that characterize nonlinear or nonmonotonic distributions, such as *α* = 1, *β* = 5 and *α* = 2, *β* = 5.

